# Systematic analysis of insertions signature in gnomAD revealed large set of novel processed pseudogenes

**DOI:** 10.64898/2026.01.15.699685

**Authors:** Artem Podvalnyi, Varvara Kucherenko, Natalia Doroschuk, Elizaveta Sarygina, Olesya Sagaydak, Olga Mityaeva, Viktor Bogdanov, Julia Krupinova, Mary Woroncow, Pavel Volchkov, Eugene Albert

## Abstract

Pseudogenes are non-functional copies of protein-coding genes that arise through genomic duplication or retrotransposition. Processed pseudogenes (PPs) is the most abundant class of pseudogenes, which is generated via mRNA reverse transcription and subsequent cDNA integration. Presence of PPs complicates the analysis of short read sequencing data due to high similarity with parental gene and frequent absence from reference genome. Here we demonstrate that the presence of non-reference (absent from reference genome) PPs leads to the very distinctive artefact of germline variant calling - long insertions on exon-intron boundaries, which sequences could be mapped to other exons of the same gene. We showed that by detecting these artifacts it is possible to identify non-reference PPs existence based on the cohort summary statistics without analysing sample-level data. We used identified signature of PPs presence to systematically mine the gnomAD database which currently contains over 70,000 whole-genome and over 700,000 exome samples to describe novel non-reference PPs. Our approach uncovered 1498 non-reference PPs of which 1268 were novel and absent in the latest GENCODE release. This resource enhances the accuracy of variant interpretation and contributes to a deeper understanding of pseudogenes diversity across human populations.

## 1 Introduction

Pseudogenes are segments of genomic DNA that share high sequence similarity with protein-coding genes but have lost their ability to encode functional proteins because of frameshifts, premature stop codons, or loss of regulatory elements (Pei et al. 2012). Among the three major classes of pseudogenes (processed, duplicated, and unitary), processed pseudogenes (PPs) are by far the most abundant. They arise when mature messenger RNA from an actively transcribed gene is reverse-transcribed and the resulting complementary DNA is re-inserted into the genome, typically by LINE-1 retrotransposition machinery (Esnault et al. 2000). Because this mechanism bypasses intron excision and promoter capture, PPs are intronless, promoter-deficient copies that nevertheless preserve much of the exon sequence of the parental gene. More than 15000 PPs have been annotated in GRCh38, and many more remain uncharacterized or absent from the reference assembly, particularly in subtelomeric and pericentromeric regions that are difficult to resolve with short reads (Ten Berk de Boer et al. 2023; Podvalnyi et al. 2025).

The near-identity between many PPs and their cognate exons poses a challenge for short-read aligners, which often lack the context needed to discriminate between paralogous loci. Therefore, reads originating from a PP can be misaligned to the parental locus. The problem becomes even more pronounced when PP is missing from the reference sequence. Such shadow mapping events propagate downstream into variant calling, transcript quantification, and structural-variant discovery, leading to spurious or mis-assigned variants (Podvalnyi et al. 2025). PP-derived artefacts have been implicated in false-positive diagnosis of pathogenic variants in genes such as PMS2, SMAD4, GBA and many others where extensive pseudogene homology confounds routine diagnostics (Millson et al. 2015; Gustavsson et al. 2024; De Vos et al. 2004).

Correct reads mapping near exon boundaries are specifically difficult for the cases of PPs absent from the reference sequence. When an intron-free PP-derived read is forced to align against the canonical gene, the aligner must introduce a large gap (representing the intron present in the gene but absent in the PP). Depending on scoring parameters, this gap may be interpreted as a cluster of mismatches or, more commonly, as a large insertion directly adjacent to the splice junction. Because such artefacts coincide with functionally critical splice sites, they are prone to misclassification as loss-of-function mutations, inflating disease burden estimates and misleading gene-essentiality analyses. Several strategies have been proposed to mitigate PP interference, including mask-based alignment to curated pseudogene databases, split-read mapping, and long-read validation (Muntéet al. 2024; Podvalnyi et al. 2025; Ten Berk de Boer et al. 2023). However, these approaches either rely on incomplete catalogues of PPs, incur substantial computational overhead, or remain inaccessible in large-scale population studies.

GnomAD (Genome Aggregation Database) offers an unparalleled resource for exploring PP-driven artefacts at scale: its current release aggregates more than 100,000 whole genomes and more than 700,000 exomes spanning diverse ancestries, providing both depth and breadth for population genetics (Chen et al. 2024). Notably, recent work suggests that at least two-thirds of individuals harbour one or more non-reference PPs, often in a population-specific manner (Ten Berk de Boer et al. 2023).

In this study we demonstrated that specific insertion signature is associated with the presence of non-reference PPs in the cohort. Moreover, we showed that by analysing only insertions on the cohort level it is possible to identify the non-reference PPs without requiring sample level information. Analysing gnomAD v4.1 we identified 14886 insertions associated with non-reference PPs. These insertions marked the presence of 1498 non-reference pseudogenes in the cohort of which 1268 are absent from the gencode PPs list. Together, these results extend the spectrum of known PPs as well as mark the systemic artifact in germline variant calling.

## 2 Results

### 2.1 Intron-Exon boundaries harbour excessive amount of germline insertions which align to the adjacent downstream exons

Previously we observed that non reference PPs presence in the sample is associated with the false positive (FP) germline variant calls in the parental gene (Podvalnyi et al. 2025). Moreover, the majority of these FP calls were insertions around splice sites. Therefore we hypothesise that by thoroughly analysing long insertions in germline variant calls we can identify the presence of PPs in the data without additional analysis of structural variants. Such an approach would allow the analysis of large summary datasets, such as gnomAD, without requiring the sample level information. To test this hypothesis, we first analysed the distribution of insertions near exon–intron boundaries in a cohort of 13307 30x coverage genomes which we previously described. Analysis of small insertions residing within ±50 bp of annotated splice sites revealed a sharp, distance-dependent enrichment that peaks within the first 5 bp from the junction (Figure 1A). Such an excess of insertions at exon–intron boundaries is consistent with the idea that reads originating from PP in a given sample are then aligned to the reference genome which lacks the corresponding PP. We focused on insertions located within ±5 bp of exon–intron boundaries and selected all of them with the length above 10bp to assure robust alignment on the next analysis step. Due to slight variations in the length of the aligned part of the read between samples, a single genomic position often harbors several insertions with nearly identical sequences. For example, the position chr2:178,450,414 in gnomAD v4.1 has 43 insertions which share the same starting sequence. We retained only one (shortest) insertion per unique genomic location. These representative insertions were subsequently realigned to the spliced MANE transcript sequences of their genes of origin. To evaluate alignment quality, we examined the extent to which each insertion sequence matched the transcript. As shown in Figure 1B, many insertions at least partially align to exonic sequences. Based on the obtained histogram, we set a threshold: an insertion was assumed to originate from a processed pseudogene if at least 30% of its sequence aligned contiguously to the cDNA without gaps. Of the 1153 insertions tested, 761 (66%) met this criterion. Of these 645 were aligned to the start of one of the exons downstream from the exon with insertion. Specifically, 347 (53.8%) matched the immediately downstream exon, and 253 (39.2%) matched the exon two positions downstream with the rest of the insertions 45 (7%) matched one of the more downstream exons. In total, supplementary table 1 lists 314 genes harbouring these 645 insertions.

**Fig. 1.**
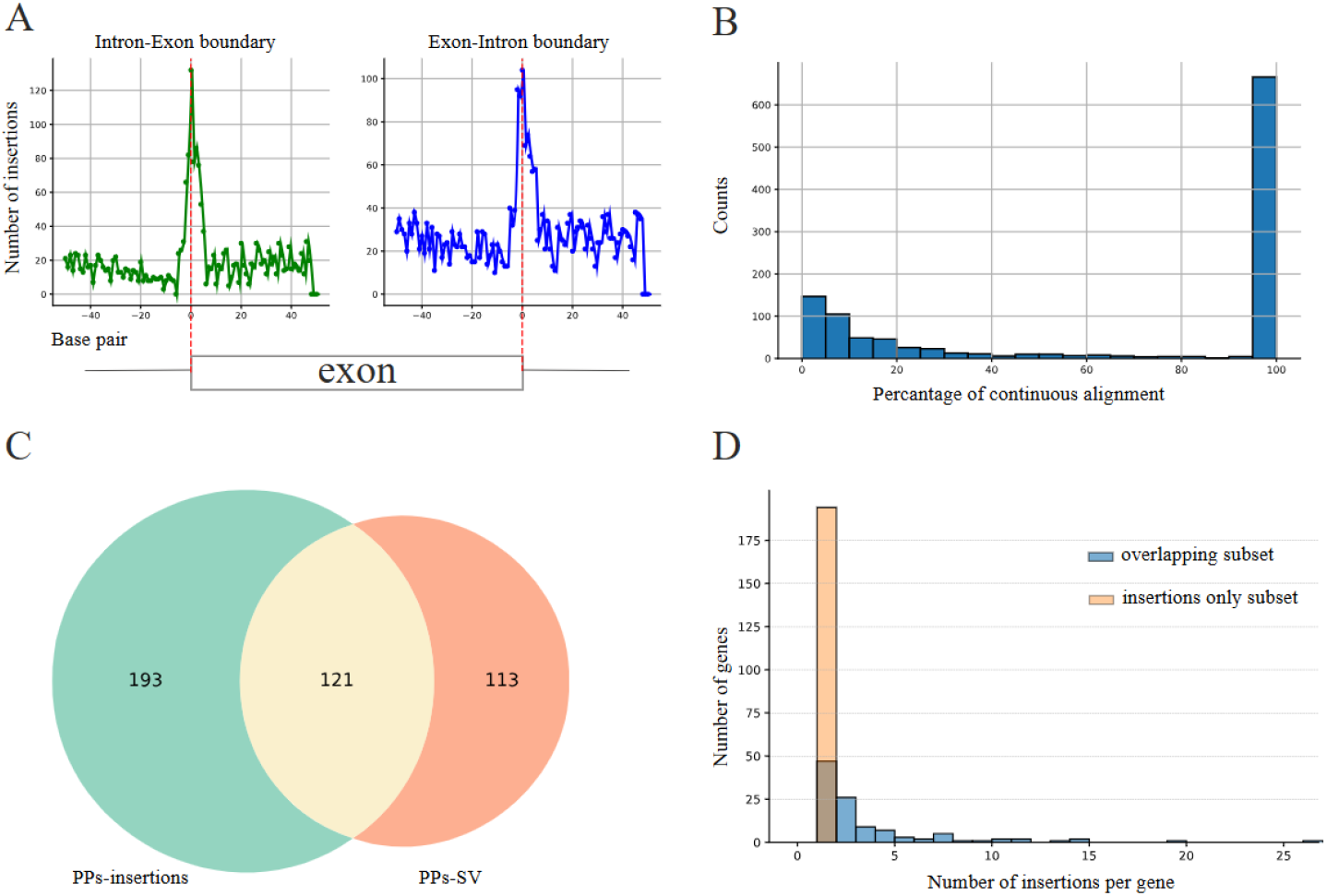
Identification of PP by analysing insertions from the internal cohort of 13307 individuals. (A) Distribution of small insertions relative to annotated exon–intron junctions of MANE transcripts in 13,307 high-coverage genomes and exomes. A pronounced peak of insertions is observed within ±5bp of the boundary. (B)Alignment identity of insertion sequences to spliced MANE transcripts. (C) Overlap between processed pseudogenes (PPs) detected by insertion-based analysis and those, identified by a structural variant approach.(D) Number of supporting insertions per PP. All PPs supported by more than two insertions belong to the validated PP-SV set, while single-insertion calls dominate the PP-insertion-only subset.

### 2.2 PPs identified by intron-exon boundaries insertions are validated by orthogonal methodology

The presence of insertions on intron-exon boundaries which aligns to downstream exons is a strong indication that a given sample might have a PP in a given gene. To test that we turned to structural variant-based approach results which we previously applied to the same cohort (Podvalnyi et al. 2025). Briefly the approach relies on detecting multiple intronic deletions within a gene as a proxy for presence of PP. For clarity, we will refer to the PPs identified by the analysis of structural variants as PP-SV, and to those identified in the present study through the analysis of insertions near exon–intron boundaries as PP-insertion. We used the PP-SV as a validation dataset as far as this approach relies on a sample level analysis of raw reads and was previously used for PP detection by our and other groups. In total our cohort has 234 PP-SVs and 314 PP-insertion. Figure 1C demonstrates the overlap of these two sets of predicted PPs. This intersection revealed a precision of 38.5% for the insertion-based catalogue (121 of 314 PP-insertions were confirmed by SVs) and a recall of 51.7% relative to the SV-based set, indicating that while the insertion-based approach captures over half of the validated PPs, it also introduces a substantial number of potential artifacts. To explore whether this noise could be filtered, we examined the number of supporting insertions per potential PP (see Fig. 1D). Strikingly, all PPs supported by two or more insertions occurred exclusively in the overlapping group, while every PP unique to the insertion-based set was supported by a single insertion. Therefore, we classify PPs with two or more supporting insertions as high-confidence, and label those supported by a single insertion as putative. Together, these observations support the possibility of identification of PPs analysing only cohort summary statistics.

### 2.3 Analysis of Gnomad revealed 1268 novel non-reference processed pseudogenes

We then applied the insertion-centric strategy developed on our cohort to the gnomAD v4.1 dataset. We scanned ±50 bp around canonical splice junctions using data from gnomAD genomes and gnomAD exomes combined. We observed the same peak of insertions around the exon-intron boundary as in our internal cohort (Figure 2A). Strikingly, we did not observe a peak on the intron-exon boundary which most likely is due to different mapper settings used in the pipeline. Aligning and filtering the set of insertions in +-5bp range from exon-intron boundary according to procedure described for the internal cohort resulted in 14886 insertions over 8729 unique positions (Supplementary table 2). Overall these insertions were distributed among 2949 candidate PPs (Supplementary table 3). Of these, 1498 PPs were supported by two or more independent insertions and, therefore, were classified as high-confidence PPs. Comparison with the current GENCODE release (v44) showed that 1268 of these high-confidence loci are not listed there as PPs. underscoring the ability of our approach to uncover previously unreported retrocopies. To further validate our identified list of 1498 identified PPs we subjected it to cell type enrichment analysis (Figure 2B). The resulting enrichment profile is dominated by several fetal cell types. Such enrichment is consistent with the known burst of retrotransposition that accompanies genome-wide demethylation in primordial germ cells providing another supporting evidence for our methodology.

**Fig. 2.**
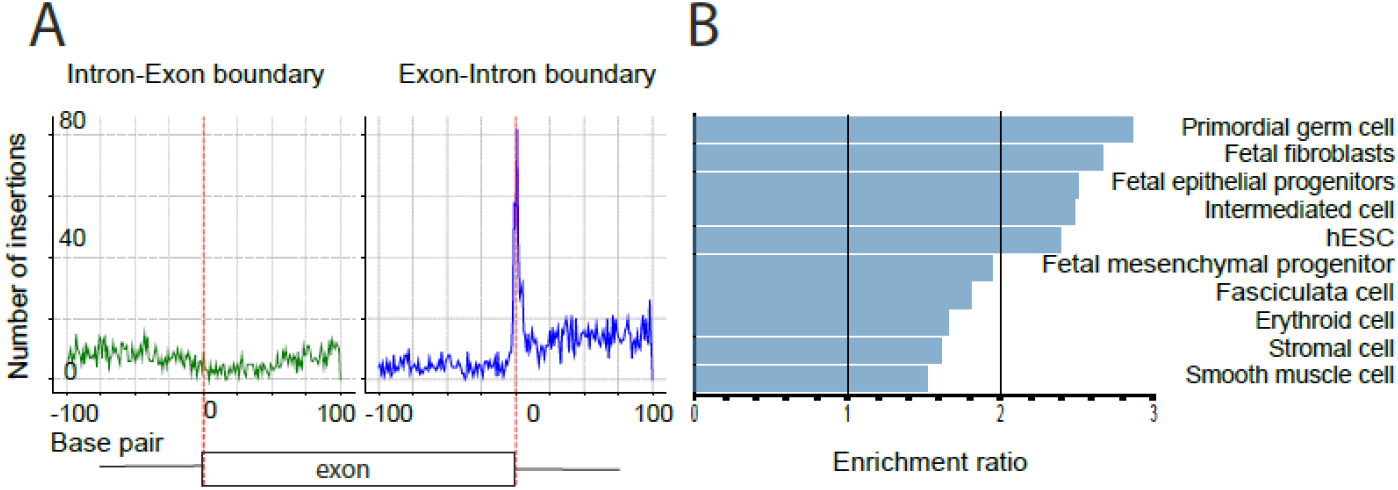
Identification and functional characterization of processed pseudogenes in gnomAD v4.1 by analysis of insertions at exon-intron boundaries. (A) Distribution of insertions relative to annotated exon–intron junctions ( MANE transcripts) for chr13 in genomes and exomes from gnomAD v4.1, plotted within ±100 bp. (B) Cell type enrichment profile of parental genes for identified PPs.

### 2.4 PPs associated insertions are mostly annotated as functionally significant due to their proximity to splice sites

To characterise variants in gnomAD which are associated with PPs and presumably are the artifacts of variant calling we revisited the set of 14886 variants. First of all, the majority of these insertions were flagged by one of the Gnomad quality filters, marking only 503 as high confidence variants (Fig. 3A). This fact supports the idea that these variants are mainly calling artefacts caused by the presence of non-reference PPs in individual genomes. Due to close proximity to exon-intron boundaries majority of these insertions are annotated as functionally significant by main gnomAD computational predictors - spliceAI (Jaganathan et al. 2019) and Pangolin (Zeng and Li 2022) for influence on splicing, PhlyoP (Pollard et al. 2010) for conservation and CADD (Rentzsch et al. 2019) for overall functionality (Fig. 3B).

**Fig. 3.**
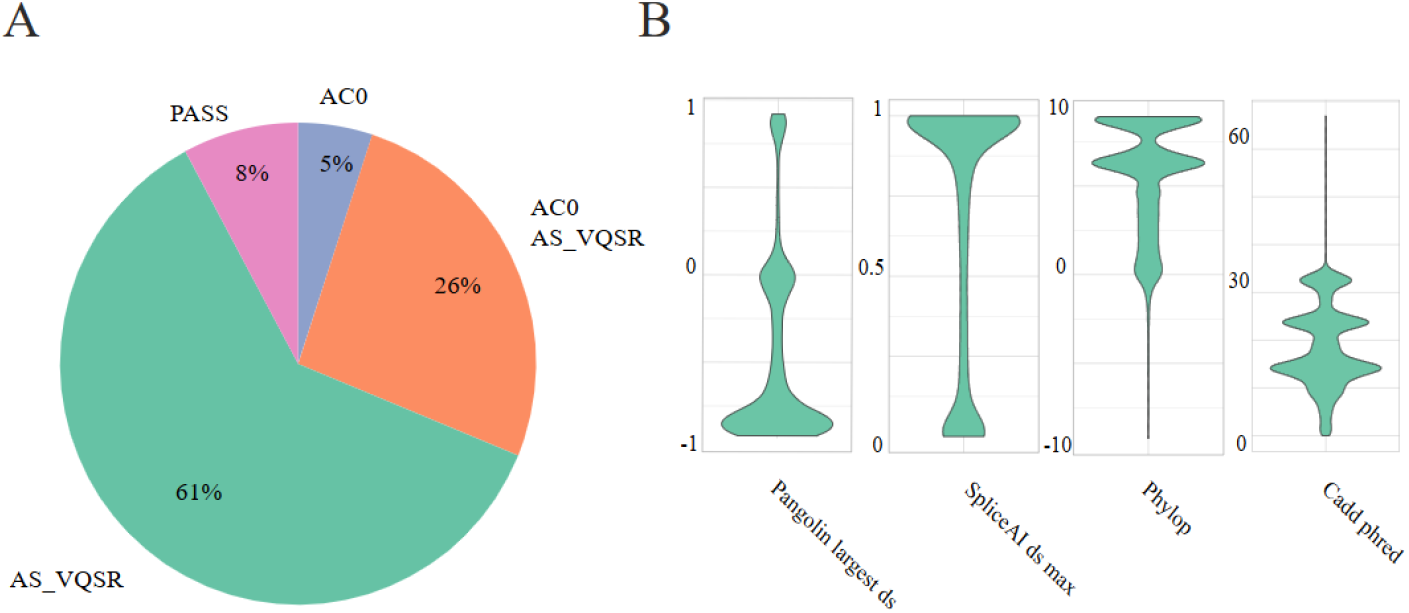
(A) Distribution of gnomAD filters among 14886 insertion alleles present at the 1498 high-confidence PP genes. (C) GnomAD functional scores distribution for these variants.

### 2.5 Identified novel PPs mostly represented by individual carriers

Because gnomAD provides variant frequencies across major human populations we also were able to describe populational distribution of identified PPs. We reanalysed a set of 14886 insertions which we previously identified as PPs associated. For each population we sum frequencies for insertions which share the same position and then average resulting frequencies for each individual gene. Clustering of the resulting z-scored gene-population matrix demonstrated that the majority of identified PPs presented in a single population (Fig4 A). Nevertheless, identified PPs can not be used as a populational marker as far as their individual frequency is low (mean = 0.0003, median 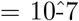 Figure 4B). Such a low frequency points out that identified PPs mostly were seen in a couple of individuals in each population.

**Fig. 4.**
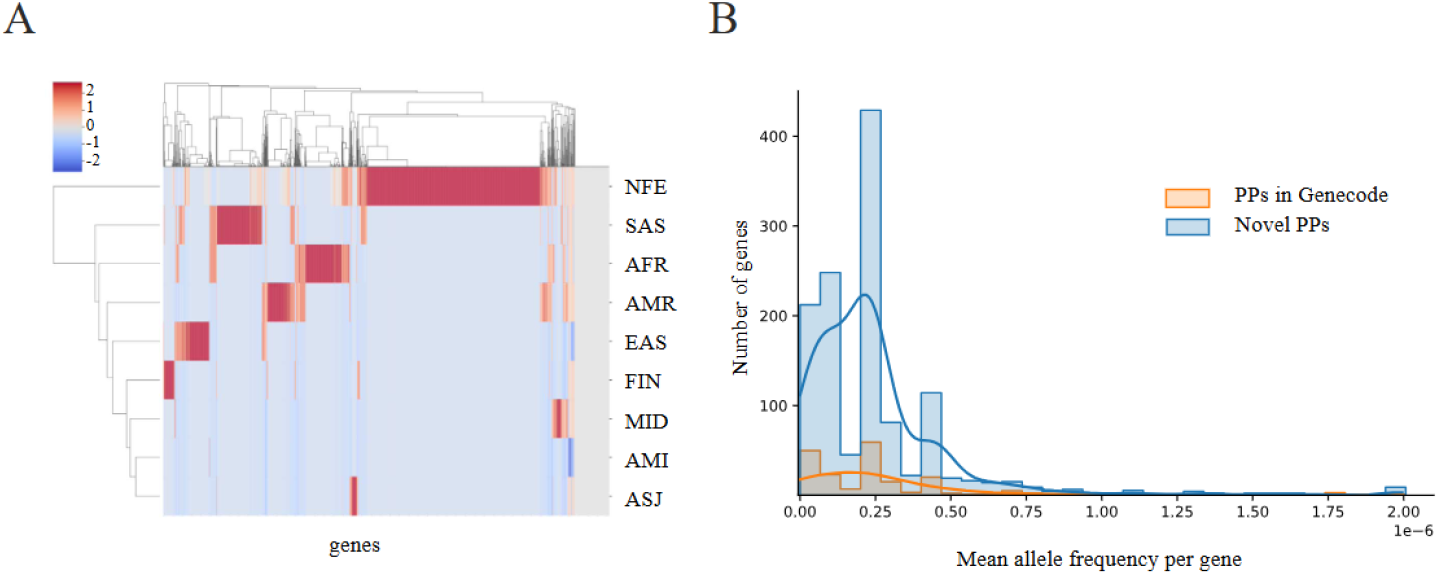
Distribution of identified PPs across main populations of gnomAD v4.1. (A) Z-scored aggregated frequencies of insertions identified at a given gene. (B) Distribution of aggregated frequencies of insertions identified at a given gene for novel PPs and PPs from Genecode.

## 3 Methods

### 3.1 Internal dataset processing and pseudogene validation list

We analyzed a cohort of 13307 standard 30x WGS samples. Samples were collected by LCC “Evogen” during routine sequencing of the healthy individuals from Russian population.. The initial analysis of the samples was performed using a modified version of the GATK Best Practices workflow. The modification included a change from BWA-MEM to minimap2 (v2.17-r941) and a change from GATK HaplotypeCaller to GATK HaplotypeCaller spark (v4.6.0.0). Previously, using that dataset, we described a pipeline for identification of PPs from the WGS data utilising the information on structural variants. Briefly, the pipeline relies on the fact that the presence of non-reference PP in WGS data produces artefacts in the structural variants calls in the form of deletions of introns of the parental gene. Here we are using the results as a validation set for developing an approach to detect PPs using only cohort summary statistics. In the current manuscript we refer to that set as PP-SV (Podvalnyi et al. 2025).

### 3.2 Identification of insertions originated from PPs

Firstly, insertions around splice sites were extracted. To identify insertions adjusted to the splice sites genomic annotations from GENCODE version 43 (Mudge et al. 2025) were used. Insertions within ±50 base pairs of annotated exon–intron junctions of MANE transcripts were extracted. The density distribution of these variants relative to splice sites was plotted to identify pattern of enrichment. Based on the obtained results (Fig. 1A) insertions within +-5 base pairs of exon–intron boundaries were selected for further analysis. The same strategy for selection of insertions was applied to the gnomAD dataset (Fig. 2A). Secondly, all insertions in splice site areas were aligned to the sequences of the processed mRNA of MANE transcript for given genes. Alignment was performed using the pairwise2 method from Biopython (Cock et al. 2009) library with the following settings: the gap opening and extension penalties were set to −10 and −4, respectively. Alignment of insertions were required to cover at least 30% (based on observed distribution, fig 2B) of the insertion length and match one of the downstream exons to be considered as PPs associated.

### 3.3 GnomAD processing

Insertions from gnomAD v4.1 were analysed according to the process described for the internal dataset. Genomes and exomes data were merged prior to analysis using bcftools v1.15 (Danecek et al. 2021). Multiallelic sites were splitted during the merge.

### 3.4 Population-Specificity Analysis

To explore the population distribution of pseudogene-derived insertions, obtained data was stratified according to gnomAD population groups, including: African (AFR), Latino/Admixed American (AMR), East Asian (EAS), Non-Finnish European (NFE), Finish (FIN), Amish (AMI), Middle Eastern (MID), Ashkenazi Jewish (ASJ) and South Asian (SAS) cohorts. Frequencies of insertion sharing the same genomic position were first summed up for each population and these numbers were then averaged across each given gene and further z-scored. That procedure resulted in a single number which characterised the mean frequency of insertions associated with splice sites for each gene with such insertions for each population.

### 3.5 Cell type Enrichment

Cell type enrichment analysis was performed using the webGestalt tool (Elizarraras et al. 2024). 1498 high confidence PPs identified from gnomAD v4.1 were subjected to over-representation analysis with functional database “cell-type” functional dataset “Human Cell Landscape”. List of “genome protein coding” genes was used as a reference dataset.

## 4 Discussion

In this study, we demonstrate that germline callsets from large cohorts harbor characteristic insertion enrichment at exon–intron boundaries. Moreover, these insertions can serve as a reliable signal for the presence of processed pseudogenes (PPs) in short-read sequencing data. By leveraging these artefactual insertions, we developed and validated an insertion-centric strategy that enables the de novo identification of non-reference PPs across large genomic datasets. Our approach overcomes key limitations of prior methods, such as the reliance on analysis of raw sample-level sequencing data for structural variants which is either computationally prohibited for large cohorts or impossible due to the lack of the access to data. The proposed approach was initially validated on a cohort of 13307 30x WGS for which we previously identified non reference PPs by analysing structural variations. Analysing only variant frequency table for the cohort we were able to detect enrichment of insertions within ±5bp of splice site. Realignment of these insertions to spliced cDNA sequences confirmed that many of them originated from downstream exons within the same gene, supporting their pseudogenic origin. Importantly, PP candidates supported by two or more insertions in a cohort exhibit full concordance with the validation set of PP identified through the analysis of structural variants based on sample-level information. This filter provides a straightforward criterion to distinguish high confidence PPs from spurious single-insertion calls, significantly enhancing the specificity of the approach. Applying this pipeline to the gnomAD v4.1 variant dataset we uncovered 2949 candidate PPs, of which 1498 met developed high-confidence criteria. Among these, 1268 loci were absent from the latest GENCODE release (v44), suggesting that a substantial number of retrocopies remain unannotated, even in current reference resources. We also identified 14886 variants from the gnomAD database, which most likely originates from PPs and therefore should be treated with caution during the analysis of individual genomes. Indeed, demonstrated distribution of gnomAD filters support the idea that identified insertions are the artifacts of variant calling. Nevertheless it has to be noted that due to close proximity of these insertions to splice sites the mostly annotated as functionally significant by in-silico predictor integrated into gnomAD. That aligns with our previous observation that the presence of non reference PPs leads to incorrect calling of clinically significant variants.

Correctness of our approach was additionally supported by the analysis of identified PPs. Cell type enrichment analysis of parental genes associated with novel PPs revealed strong overrepresentation of cell types from early development with primordial germ cells being the top hit. This aligns with previous observations that retrotransposition is especially active in the context of genome-wide demethylation, particularly in primordial germ cells. The transcriptomic surge during this developmental window likely fuels the retrotransposon machinery with abundant mRNA templates, giving rise to germline-encoded retrocopies.

From a population genetics perspective, although preliminary inspection suggested potential population-specific pseudogene patterns, deeper analysis revealed that most of these cases were driven by single-individual events.

The clear limitation of the proposed methodology for identification of non reference processed pseudogenes from the cohort summary statistics is a low sensitivity. Analysing summary statistics of the internal cohort of 13307 individuals for which non-reference PPs were identified through the analysis of the structural variants we were able to uncover only 121 out of 234 PPs. That points out that despite the 1498 identified here non-reference PPs in gnomAD, the analysis of a raw sequencing data would uncover substantially more variability in individuals genomes associated with PPs.

## 5 Conclusion

In conclusion, our study presents a robust, alignment-based framework for the detection of non-reference processed pseudogenes at scale from summary statistic data, by exploiting characteristic insertions artifacts rather than relying on long-read sequencing or assembly-based annotation. The resulting list high-confidence PPs complements existing annotations and improves variant-calling fidelity.

## Supplementary information

- Supplementary table 1 - genes for which splice-site associated insertions were found in the internal cohort
- Supplementary table 2 - PPs associated insertions from gnomAD v4.1
- Supplementary table 3 - Genes with splice-site associated insertions identified in gnomAD
- Supplementary table 4 -

## Declarations

- Funding The research was supported by the Ministry of Science and Higher Education of the Russian Federation (agreement # № 075-03-2024-117/6, project # FSMG-2024-0029) (to Viktor Bogdanov) and Russian Science Foundation (Grant № 23-64-00002) (to Olga Mityaeva and Pavel Volchkov)
- Conflict of interest/Competing interests ND, ES, OS were employed by LCC Evogen. The remaining authors declare no conflict of interest.
- Ethics approval and consent to participate The study was approved by the local ethical committee of the Independent Multidisciplinary Committee on Ethical Review for Clinical Trials (Moscow, Russia) and was performed in accordance with the approved guidelines and the Declaration of Helsinki. Inform consent was obtained from all participants.
- Consent for publication
- Data availability
- Materials availability
- Code availability
- Author contribution

